# Logic-based Analysis of Gene Expression Data Predicts Pathway Crosstalk between TNF, TGFB1 and EGF in Basal-like Breast Cancer

**DOI:** 10.1101/614933

**Authors:** Kyuri Jo, Beatriz Santos Buitrago, Minsu Kim, Sungmin Rhee, Carolyn Talcott, Sun Kim

**Author notes:** Equal contributors (Kyuri Jo, Beatriz Santos Buitrago).

## Abstract

For breast cancer, clinically important subtypes are well characterised at the molecular level in terms of gene expression profiles. In addition, signaling pathways in breast cancer have been extensively studied as therapeutic targets due to their roles in tumor growth and metastasis. However, it is challenging to put signaling pathways and gene expression profiles together to characterise biological mechanisms of breast cancer subtypes since many signaling events result from post-translational modifications, rather than gene expression differences.

We present a logic-based approach to explain the differences in gene expression profiles among breast cancer subtypes using Pathway Logic and transcriptional network information. Pathway Logic is a rewriting-logic-based formal system for modeling biological pathways including post-translational modifications. Proposed method demonstrated its utility by constructing subtype-specific path from key receptors (TNFR, TGFBR1 and EGFR) to key transcription factor (TF) regulators (RELA, ATF2, SMAD3 and ELK1) and identifying potential pathway crosstalk via TFs in basal-specific paths, which could provide a novel insight on aggressive breast cancer subtypes.

**Availability:** Analysis result is available at http://epigenomics.snu.ac.kr/PL/

## 1 Introduction

Breast cancer is the first cause of cancer death in women with a worldwide mortality rate of 12.9% [1]. A better understanding of the pathogenesis of the disease is a key to improving current therapies and discovering new treatment options. Especially, signaling pathways in breast cancer such as tumor necrosis factor (TNF), transforming growth factor beta 1 (TGFB1) and epidermal growth factor (EGF) have been studied as therapeutic targets due to their roles in tumor growth and metastasis [2, 3, 4]. On the other hand, genome-wide gene expression profiles are measured for many breast cancer patients using gene expression measurement technologies such as gene expression microarray and sequencing machines. As a result, clinically important breast subtypes are well characterised at the molecular level in terms of gene expression profiles [5, 6]. However, there is little study in combining signaling pathways and gene expression profiles together to characterise biological mechanisms in each of breast cancer subtypes since many signaling events result from post-translational modifications (PTMs) followed by gene expression differences [7], which is difficult to be identified using the existing pathway models. The goal of this study is to provide an integrated analysis of signaling pathways and gene expression profiles to determine subtype-specific path from key receptors to key transcription factor (TF) regulators to downstream genes targeted by TF. There are several challenges for achieving this research goal.

One approach to model cancer phenotype is to construct *in silico* gene regulatory networks (GRNs) based on the putative relationship (e.g. activation or inhibition) between genes inferred from gene expression data. Network inference algorithms like ARACNE [8] have been used to elucidate the underlying mechanisms of breast cancer such as Epithelial-Mesenchymal Transition (EMT) [9] and metastasis [10]. As breast cancer subtypes can be defined at the molecular level using gene expression information [5], these network-based machine learning approaches are very powerful to bring novel insight into breast cancer biological mechanisms. However, *in silico* network generated from gene expression without prior knowledge requires additional analysis steps due to a number of false positive relationships.

To overcome the limitation of gene expression data, several algorithms have incorporated the information of biological pathways into the analysis in addition to gene expression data [11]. Such pathway analysis algorithms use networks generated from the curated knowledge of biological pathways provided by public databases such as KEGG [12] and NCI PID [13], rather than the network generated by *in silico* prediction. Thus, they charts a more precise and compact landscape of dysregulated biological processes, suitable for comparing networks from multiple phenotypes. When converting the information from the public databases into a network, however, most of the pathway analysis tools such as Clipper [14] construct a simple gene-gene network without PTM reactions (e.g. phosphorylation or ubiquitination), leading to a loss of information on pathways [15]. Simplified network provides a limited view of cancer subtypes since many biological events such as signal transduction can occur without significant changes in expression levels. For instance, a signaling pathway includes a sequence of PTM reactions that can modify the activity of TF proteins prior to their transcriptional regulation on the target genes [16].

Logic modeling is another approach to model and analyse a complex pathway. In such models, molecular events including PTMs are defined as general rules and a set of rules makes up a knowledge base. By applying rules to the premise (initial state of the cell), the user can derive a conclusion (final state of the cell). Several logic-based models have been developed so far [17], especially boolean models with two binary states for floral morphogenesis [18], mammallian cell cycle [19], apoptosis [20] and hepatocyte growth factor (HGF) [21]. Recently, BioASF [22] suggested a general framework for pathway models specified in BioPAX language [23] that has been used by several pathway databases [24, 25]. Although these models delineate each biological process in more detail than a gene-gene network used in the pathway analysis, the main purpose of the logic models is the simulation study to enumerate possible final states or concentrations of the product, not designed for integrating gene expression data and performing comparative studies between phenotypes.

Pathway Logic (PL) [26] is a general system for modeling signaling and metabolic pathways in Maude language [27]. PL provides several knowledge bases including STM7 knowledge base with 32 models of signal transduction for different stimuli. Models in PL are constructed with detailed descriptions of biological process including cellular locations, interactions and PTMs of the molecules. Unlike the other logic models, PL system provides internally implemented functions such as comparison between two different models and dynamic searching of subgraph with certain endpoint conditions, showing a wide range of application for the pathway analysis beyond the typical simulation studies. Recently, phosphoproteomics data has been integrated into the PL system to derive different response of signaling pathways to different drugs in SKMEL133 melanoma cancer cells [28]. This research work shows that integrating the PL system and omics data can effectively narrow the range of rules in deductive reasoning by specifying the condition-specific cell states observed in the experiments. In fact, gene expression data has much more potential for the integrated system as millions of omics data sets have been generated from high-throughput experiments over the past decade [29].

A possible limitation of the pathway analysis might be the small set of genes belonging to one pathway (e.g. 29 to 351 genes in KEGG signaling pathways), not enough to analyse the global changes triggered by the signal transduction. One of the methods to expand the pathway analysis of a limited set of genes is to incorporate TFs. TFs are principal regulators of gene expression in eukaryotic cells [30, 31]. TFs regulate their target genes to mobilize the appropriate protein response and thereby contribute to a global change in cellular states, even resulting in a different phenotype of an organism [32, 31]. Most of the signaling pathways described in the database start from a ligand binding to a receptor and the transmitted signal activates a number of proteins, including TFs. Therefore, TFs can work as a bridge from signal transduction to broader changes in a cell or an organism.

In this study, we propose a novel computational framework to integrate pathway models of PL, gene expression data and transcriptional network to explore subtype-specific biological mechanisms in breast cancer. By rewriting rules in PL models, we derived paths of signal transduction from key receptors to key TF regulators and to downstream genes targeted by TFs. TFs included in the signaling paths are statistically tested based on their contribution to the subtype-specific expression of their target genes. Key features of the proposed method compared to existing methods are as follows:

1. Pathway models defined in PL include detailed information on biological entities such as PTMs that are often ignored in *in silico* network or current pathway network structure with limited interaction types.

2. Logical inference-based functions implemented in PL can be used for pathway analysis such as searching for sub-pathways satisfying a specific condition and also for comparing multiple pathways, which are not available in the existing logic models designed for simulation studies.

3. Integration of gene expression data with PL models and PL functions provide a more accurate and comprehensive landscape of signaling pathways —e.g. signaling paths activated only in a certain phenotype (subtype) including the upstream PTM reactions.

We used our analysis framework to analyse The Cancer Genome Atlas (TCGA) breast cancer data [33] and determined subtype-specific signaling path from three receptors (TNFR, TGFBR1, EGFR) to four TFs (RELA, ATF2, SMAD3, ELK1) and identified their roles in breast cancer. Performance of our method is compared with a reverse engineering algorithm, ARACNE [8] and a pathway analysis algorithm, Clipper [14].

## 2 Methods

The computational framework proposed in this paper is based on the pathway models in PL. In PL models, steps of a signal transduction pathway are represented as occurrences and transition rules. An **occurrence** is a protein together with its state and location. Transition **rules**, representing biochemical reactions that change a state and/or location of the occurrences, are curated from the literature (**datum**) and collected into a knowledge base. In the context of such a knowledge base, a model is defined by specifying the set of occurrences initially present (called an initial state or **dish**). The graphical representation of occurrences, rules and dishes is shown in Figure 1 (a).

**Figure 1:**
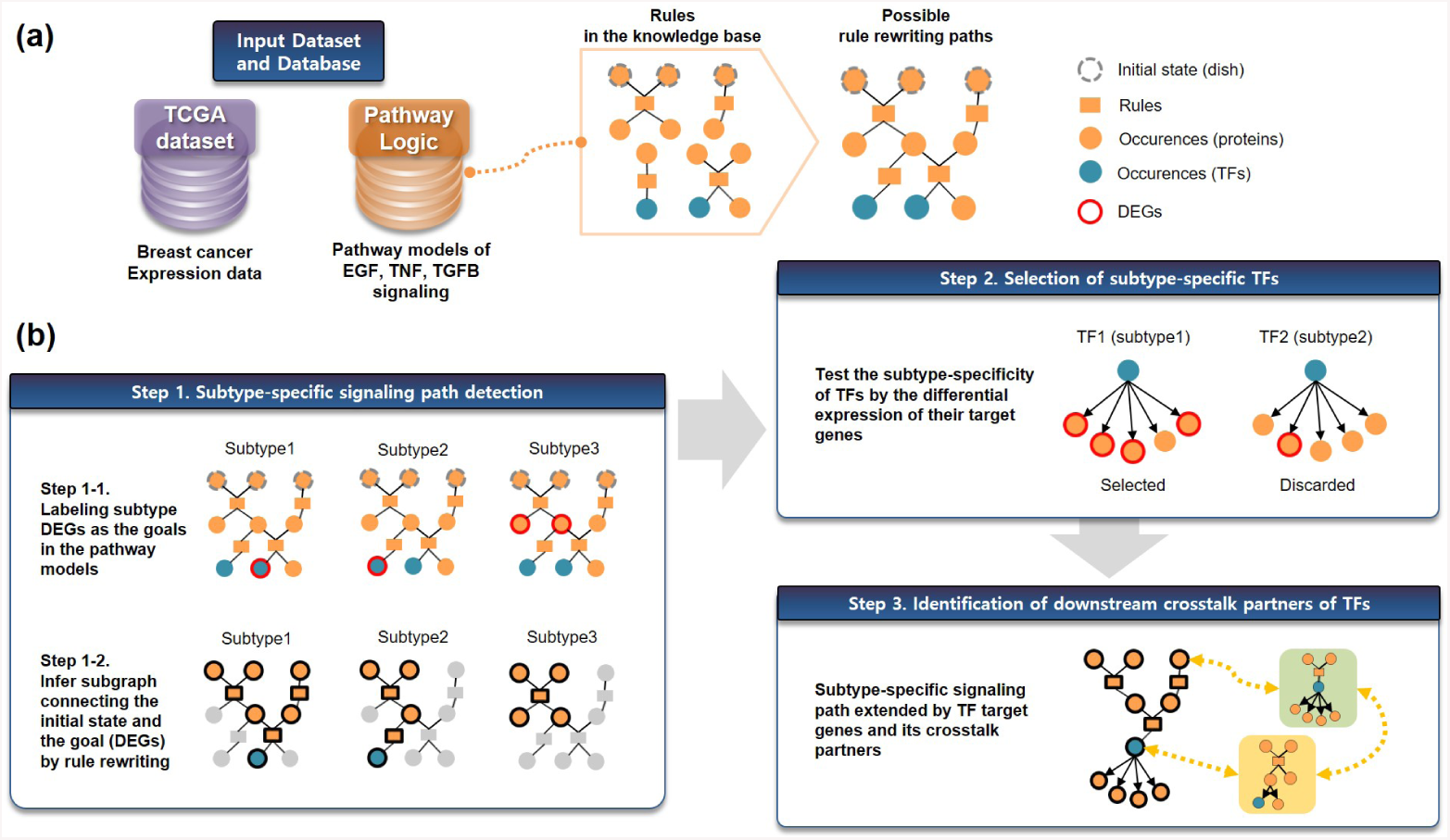
Workflow of the proposed method. (a) Two input dataset (database) for the analysis. Gene expression data was obtained from TCGA data portal [33] to analyze breast cancer subtypes. We used PL database to model signaling pathways, where molecular and cellular processes are represented with occurrences (orange circles) and rules (orange boxes). Some of the proteins in the occurrences are TFs (blue circles). (b) Three-step procedure for the analysis. To detect the subtype-specific signaling paths in breast cancer, differentially expressed genes (DEGs) were determined for each subtype and set as the goals of the rule rewriting (Step 1-1). By performing rewriting of rules in the model, we inferred subgraph of the signaling pathway starting from the initial state leading to DEGs (Step 1-2). The subgraphs include TFs, the subtype-specificity of which was tested by the differential expression of their target genes (Step 2). As a result, subtype-specific signaling paths were extended to the downstream target genes of TFs and their crosstalk partners.

Among the knowledge bases in PL, STM7 is a model of intracellular signal transduction that responses to peptides, chemicals, or stresses added to cells. STM7 includes 32 dishes (initial states), each representing one stimuli. To analyse breast cancer dataset, we selected three dishes from STM7 knowledge base according to the relevance to cancer mechanism —response to Tumor Necrosis Factor (TNF), Transforming Growth Factor Beta 1 (TGFB1) and Epidermal Growth Factor (EGF) stimulation. TNF, TGFB and EGF signaling pathways are known to have strong relationship through pathway crosstalk in breast cancer cell line modulating the tumor phenotype [34]. Table 1 shows the brief description of these PL models. Initial states and representative rewrite rules (receptor binding event of each ligand) of three pathways are described in Supplementary Information.

**Table 1:** Description of EGF, TNF and TGFB1 models in PL.

| Dish | Ligand(s) | Receptor(s) | # of Rules | # of Datums |
| --- | --- | --- | --- | --- |
| TnfDish | Tnf | TnfR1 | 87 | 1,915 |
| Tgfb1Dish | Tgfb1 | TgfbR1, TgfbR2 | 57 | 968 |
| EgfDish | Egf | Egfr | 129 | 2,281 |

### 2.1 Finding transcription factors mediating subtype-specific control of pathway crosstalk

It is well known that pathway crosstalk frequently occurs between signaling pathways to orchestrate cooperative response to various environmental stress [35]. The way two pathways communicate determines how cancer tissue reacts to oncogenic environment, which affects its potential malignancy. For example, Dent et al. revealed that a combination of pathways, including AKT-ERK crosstalk, affected cancer cell viability and had bad influence on prognosis [36].

TFs have been considered as important participants for regulating pathway crosstalk and also potential molecular targets. Blokzijl et al. suggested that TF can act as a mediator between different signaling pathways [37]. Such crosstalk between signaling pathways can be targeted by a molecular therapy if we can understand the underlying mechanism and specify targets [38]. The proposed algorithm can contribute in this regard since it finds TFs having potential as pathway crosstalk mediators and shows how the crosstalk is regulated. The analysis starts with signaling upstream factors (e.g. TNF, TGFB1, and EGF) and finds downstream paths leading to the pathway crosstalk mediated by a TF. This process can be summarized as three steps as described in Figure 1 (b).

- Step 1: Detecting subtype-specific paths from signaling upstream factors to potential TF mediators.
- Step 2: Selecting TFs that regulate subtype-specific expression of target genes.
- Step 3: Identifying downstream crosstalk partners mediated by TFs.

### 2.2 Subtype-specific signaling path detection by rule rewriting

Given that differentially expressed genes (DEGs) for each subtype of cancer are determined by statistical algorithms such as Limma [39], subtype-specific signaling path is defined as the path where the initial state of the pathway is rewritten to the state with activated subtype DEGs. Therefore, subtype DEGs up-regulated in cancer cells (tagged with `-act` modifier in PL models) are used as goals of the signaling in the PL model (Step 1-1 in Figure 1 (b))). For the four breast cancer subtypes, we had different sets of DEGs in comparison with the same normal samples (normal vs. luminal A, luminal B, basal-like or HER2-enriched) deriving different subtype-specific signaling paths.

In PL models, finding a path from an initial state to the goals (endpoints) can be performed in two ways: using the Maude language and the Pathway Logic Assistant (PLA). The Maude language provides the `search` command (Listing 1) that takes parameters (`[n]` : the first n solutions; `=>+:` at least one step) and target state of the endpoint defined by the operator `PD`. The argument of `PD` is the locations with their respective contents. Listing 1 shows an example of the search command to determine a path from EgfDish to the activated Erks protein. The contents of each location (e.g. `EgfRC`) are elements or variables (e.g. `thEgfRC:Things`). In this example, the target state of the endpoint is a protein `prot:BProtein` activated (`-act`) in the nucleus (`NUc`) with other modifications `mod:ModSet` and the variable `prot:BProtein` should belong to the `ErkS` protein family. The solution to this query is the matching state of the endpoint (Listing 2). Note that the variable `prot:BProtein` is matched to `Erks` and has modification of `phos (TEY)` and `phos (SPS)` in addition to activation. Asking Maude to `show path label [state ID]`, we can summarize rewrite rules applied to reach the state [40].

**Listing 1:**
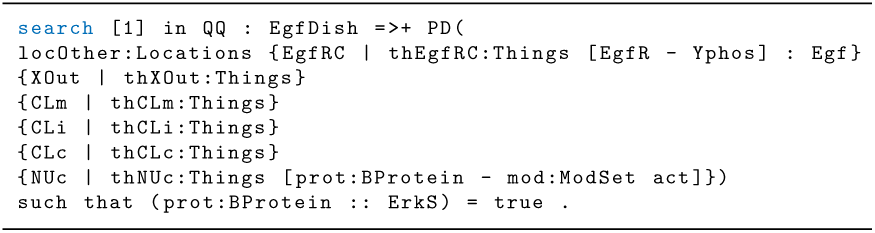
Example of search command in PL using Maude language. The code is to find a path from EgfDish to activation of Erks.

**Listing 2:**
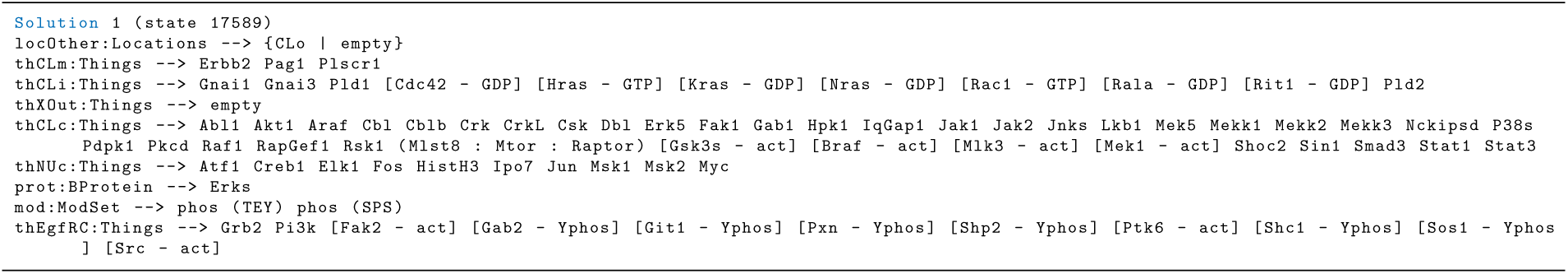
Example of solution (matching state) for search command in PL. Erks is activated and phosphorylated at two different sites.

Another approach to derive subgraph in PL is the PLA. PLA can be executed online (http://pl.csl.sri.com/) and provides an interactive graphical interface to browse and query PL network models. Using PLA, we define endpoints as checking the ‘make occ as goal’ box for the target occurrences and created the subgraph by ‘Subnet’ button. In this study, we explored three dishes —TNF, TGFB1 and EGF where the goals were occurrences of DEGs in each breast cancer subtype and computed subgraphs that led to the activation of those genes. As a result, we obtained four different subgraphs for each dish, each subgraph for each breast cancer subtype (Step 1-2 in Figure 1 (b)).

### 2.3 Selection of subtype-specific transcription factors

Each subtype-specific signaling path may include TFs that may cause a global gene expression change in the cancer subtype samples. To ensure the significance of TFs as subtype-specific regulators, we test the differential expression of their target genes ‘between subtypes’ (e.g. basal-like vs. three other subtypes of breast cancer). We assume that if a transcription factor is a subtype-specific regulator, a ratio of differentially expressed target genes to all target genes should be much higher than a ratio of all DEGs to all genes in total. A hypergeometric test was used to measure the influence of TF comparing a ratio of total DEGs (*n*) to total genes (*N*) with a ratio of DEGs among the target genes (*k*) to total target genes of the TF (*K*) (Eq. 1). P-value was calculated as one minus cumulative distribution function of Eq. 1. The hypergeometric test is a simple and fundamental metric that has been used for finding pathways with a high proportion of DEGs [41].

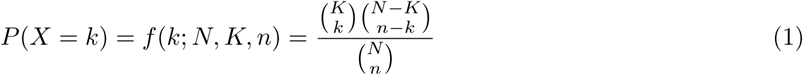

Interaction information between TFs and their target genes in a GRN was from HTRIdb (The Human Transcriptional Regulation Interactions database) [42] that provides experimentally verified or computationally predicted TF-DNA-binding sites from six public databases and literature [43]. Additionally, we used MSigDB (Molecular Signatures Database) [44] where C3-TFT gene set (gene set of transcription factor targets) is provided [45].

### 2.4 Identification of the downstream crosstalk partners of TFs

For each TF detected as subtype-specific in Step 2, we identify the downstream crosstalk partners using their target genes. Among the target genes, we filter subtype DEGs that are up/down-regulated in the same subtype where TF is activated and their enrichment on biological pathways is tested using Enrichr [46] web service available at http://amp.pharm.mssm.edu/Enrichr/. We assume that pathways enriched with TF target genes are potential crosstalk partners of TFs, deriving a subtype-specific signaling circuit connecting the upstream pathway of TF and downstream crosstalk pathways.

## 3 Results

To identify the subtype-specific signaling paths and their crosstalk in breast cancer, RNA-seq data was downloaded from TCGA data portal [33]. Differentially expressed genes (DEGs) of four breast cancer subtypes (luminal A, luminal B, HER2-enriched and basal-like) compared to normal cells are determined by voom function [47] in Limma R package [39] with a threshold of adjusted P-value ¡ 0.05. Four sets of DEGs were labeled as goals (endpoints) for rule rewriting. The DEGs had few (1 to 3) overlap between subtypes (Supplementary Fig. S1-S3), thereby generating distinctive signaling paths to the DEGs for each subtype (Supplementary Fig. S4-S6 or http://epigenomics.snu.ac.kr/PL/).

### 3.1 Subtype-specific signaling paths and transcription factors of breast cancer

From TCGA breast cancer data, we found seven different subtype-specific paths originated from three signaling factors (TNF, TGFB1, and EGF) leading to a TF (Table 2). Based on the expression pattern of target genes, we selected four TFs (red texts in Table 2) by testing the subtype-specificity. These TFs (RELA, ATF2, SMAD3, and ELK1) have target genes differentially expressed in the same subtype as their upstream signaling paths. For example, in TNF-RELA case, we found that the ratio of basal-specific DEGs are significantly larger in genes targeted by RELA than non-targeted genes. It shows that RELA regulates downstream target genes in basal-specific manner as well as RELA itself is activated by the basal-specific upstream signal from TNF.

**Table 2:** Subtype-specific signaling paths and transcription factors of breast cancer detected from PL analysis. Three (ELK1, RELA, ATF2) and one (SMAD3) TFs are detected as subtype-specific regulators for basal-like and luminal B subtypes, respectively.

| Pathway | TF | Subtype | # of DEGs | # of total genes | # of DE target genes | # of target genes | Hypergeometric P-value |
| --- | --- | --- | --- | --- | --- | --- | --- |
| EGF | **ELK1** | Basal-like | 15,676 | 20,501 | 1,488 | 1,886 | **0.0038** |
| TNF | **RELA** | Basal-like | 15,676 | 20,501 | 387 | 484 | **0.0278** |
| TGFB1 | **ATF2** | Basal-like | 15,676 | 20,501 | 38 | 43 | **0.0152** |
| TGFB1 | SMAD3 | Basal-like | 15,676 | 20,501 | 477 | 626 | 0.5476 |
| TGFB1 | **SMAD3** | Luminal B | 13,149 | 20,501 | 421 | 626 | **0.0445** |
| TGFB1 | SMAD4 | Luminal A | 13,599 | 20,501 | 428 | 637 | 0.3071 |
| TGFB1 | SMAD4 | Luminal B | 13,149 | 20,501 | 423 | 637 | 0.1045 |

Among the four TFs, RELA, ATF2 and ELK1 are all activated in the basal-like subtype but in the different signaling pathway. Those three paths all started from the common rules of all subtypes but branch off to the basal-specific rules later, mostly through the rules shared with the luminal B subtypes in the midway (Figure 2). Specific rules in each path are shown in Supplementary Figures S7-S10.

**Figure 2:**
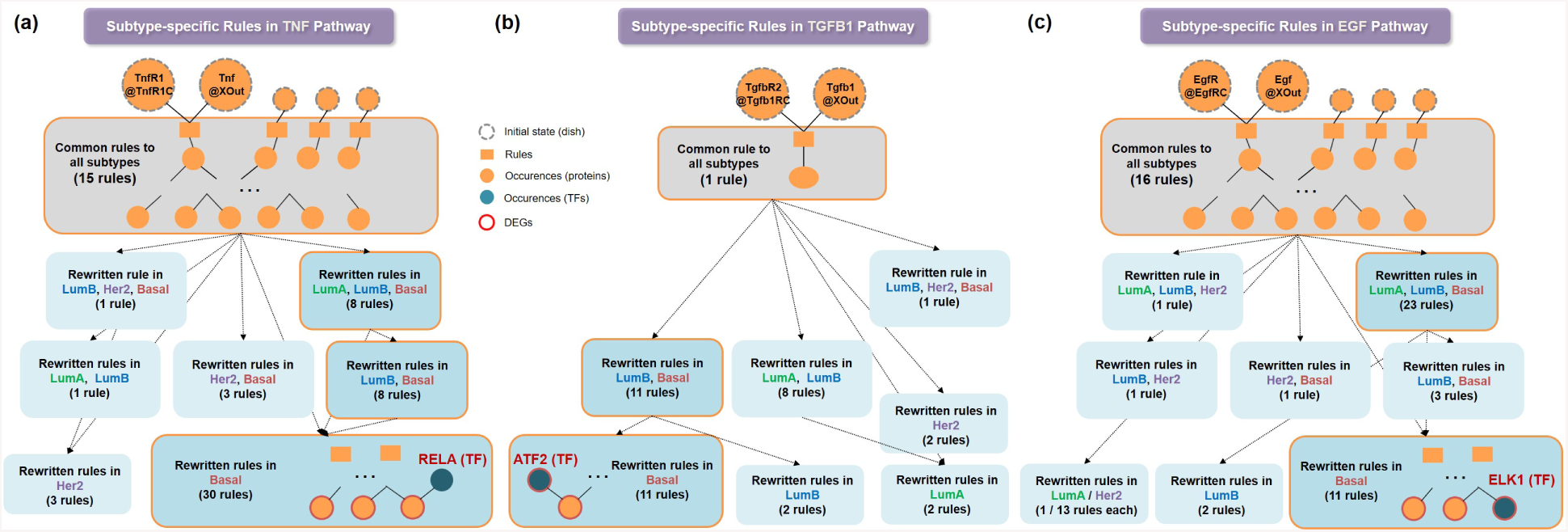
Subtype-specific signaling paths and transcription factors of breast cancer. (a) Overview of the rewritten rules in (a) TNF, (b) TGFB1, (c) EGF pathways. Transcription factors RELA, ATF2 and ELK1 are found in the basal-like-specific path of the pathways, respectively.

### 3.2 A signaling circuit connecting TNF, TGFB1, and EGF pathways specifically regulated in basal-like breast cancer

From the enrichment test on four TFs (Supplementary Table S1), we identified potential crosstalk partners of EGF, TNF and TGFB1 pathways. For example, in TNF-RELA case, 387 differentially expressed target genes of RELA are significantly enriched with five signaling pathways other than TNF pathway itself, which means that RELA can be considered as a mediator of basal-like specific communication between TNF-pathway and these five external pathways. As a result of the downstream analysis, a potential crosstalk of three pathways (TNF, TGFB1, EGF) by three mediators (RELA, ATF2, ELK1) in the basal-like subtype is discovered and summarized into a signaling circuit (Figure 3). Since a luminal B-specific path of SMAD3 is not involved in the circuit, biological meaning of the path is described in the Supplementary Information.

**Figure 3:**
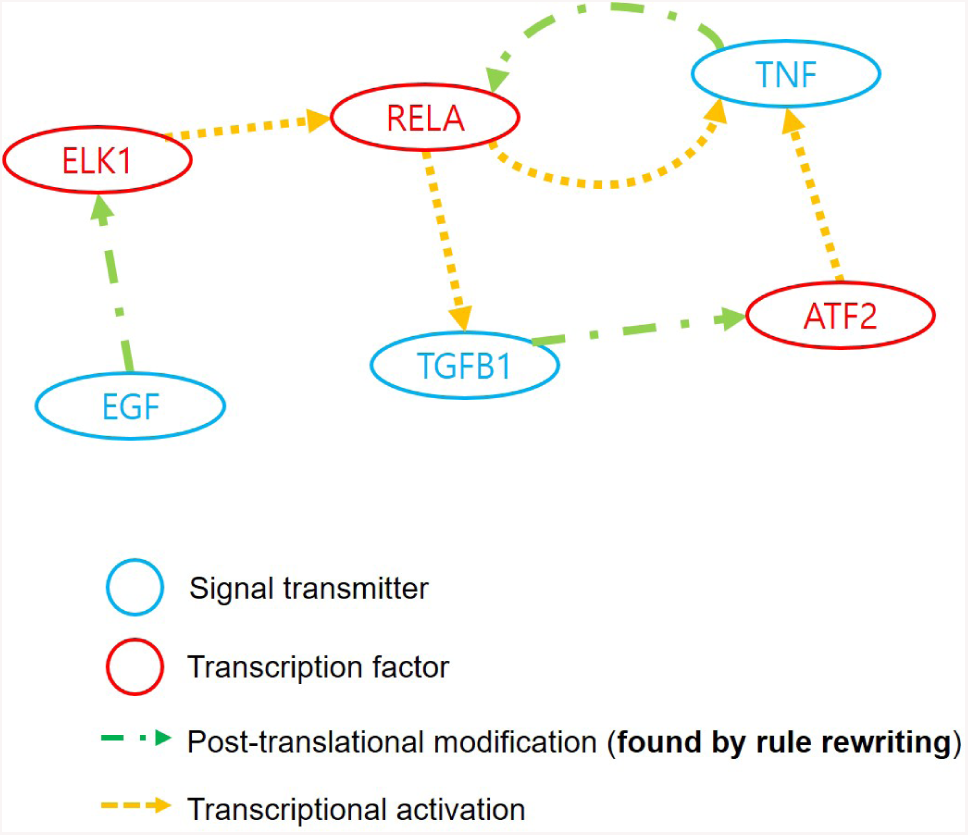
A signaling circuit connecting TNF, TGFB1, and EGF pathways. Three signaling pathways (TNF, TGFB1, and EGF) are connected to each other mediated by three TFs (RELA, ATF2, and ELK1). Blue nodes indicate upstream signaling factor genes, while red nodes indicate TF mediator genes. Green and yellow arrows represents post-translational modification (PTM) and transcriptional regulation, respectively.

In the signaling circuit derived for basal-like subtype (Figure 3), we found that three signaling factors (TNF, TGFB1, and EGF) are destined to each other as well as to external pathways. For example, the signal originated from EGF factor comes down to ELK1, then ELK1 broadcasts it to various downstream target genes, one of which is RELA. In turn, RELA propagates the signal to downstream target genes including TGFB1, which subsequently activates ATF2. Finally, ATF2 carries signal to TNF that again activates RELA. This recursively linked signaling cascade constitutes a closed circuit specifically activated in basal-like breast cancers.

Several studies have found that cooperation among the above three pathways induced malignant cancer phenotypes. For example, according to Kali et al., TNF and TGFB1 are found to synergistically increase the cancer stem cell properties in pancreatic cancer cell line [48]. Dunfield et al. also suggest that EGF inhibits the anti-proliferative effect of TGFB1, contributing to increased proliferation in ovarian cancer cells [49]. Our system reproduced the close cooperation of three pathways as known in the literature. In addition, our analysis newly discovers three TF mediators as potential links between them.

### 3.3 Systematical validation of the basal-specific signaling circuit

The signaling circuit of TNF, TGFB1, and EGF pathways is highly associated with basal-like breast cancers, thus we hypothesized that this signaling circuit is a major feature that characterising the basal-like breast cancer. To support the hypothesis, we performed two experiments:

1. A decision tree model was built with the expression values of six genes that belong to the signaling circuit to identify variables characterizing basal-like subtype.

2. For clinical implication, survival analysis was performed using TNF and ELK1 genes that were major discriminatory variables in the decision tree model.

### 3.3.1 A decision tree model for basal-risk

We built a classification model using six genes in the signaling circuit as input variables and subtype labels (basal/non-basal) of each sample as corresponding target variables. The decision tree (DT) model would show how the combination of six genes contributes to characterising basal-like subtype. To build the model, we used J48 decision tree algorithm implemented in WEKA v3.8 [50] with the confidence factor as 0.25 (default) and the minimum number of objects per leaf as 51, which is approximately 5% of total number of samples (i.e. 1,016 samples).

The primary goal of the DT model is to classify breast cancer samples (1,016 primary tumors in TCGA-BRCA) into several groups having different risk of basal-like subtype. We consider the ratio of samples having basal-like breast cancer in a group of samples as a risk and refer it as *basal-risk*. The result is illustrated in Figure 4. The model was built with gene expression profile normalized with percentile ranked in ascending order among 1,016 samples. The gene expression value of each gene was represented as percentile, where the notation *P*_*i*%_ indicates that the expression level of sample is higher than *i*% of 1,016 samples.

**Figure 4:**
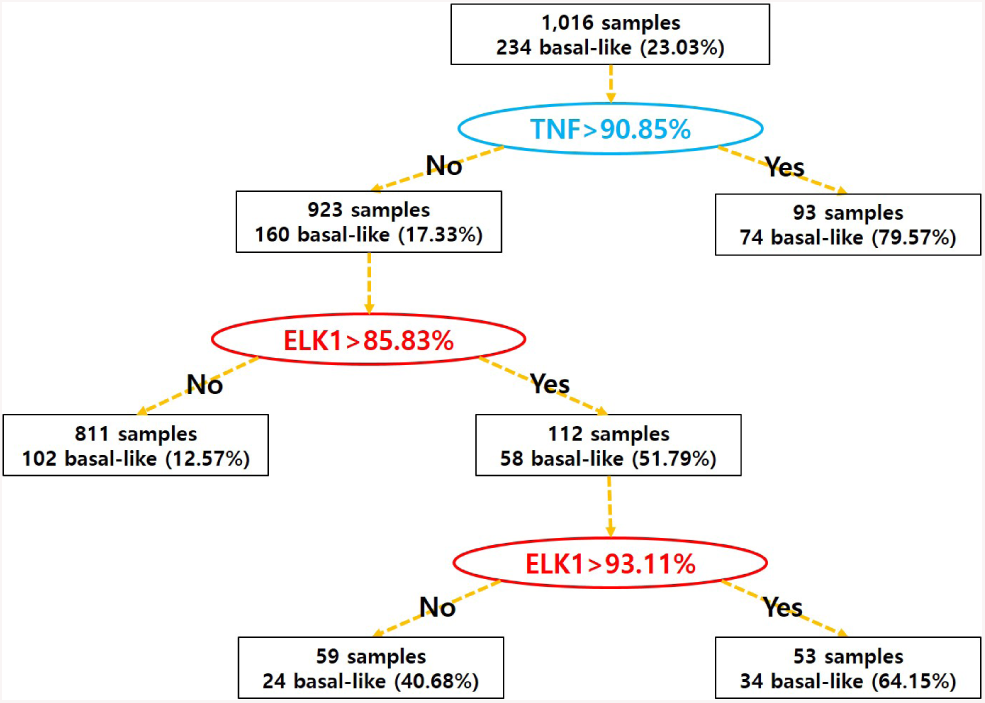
Decision Tree model constructed with six genes in the signaling circuit of basal-like subtype. 1,016 samples from TCGA breast cancer patients are used as gene expression profiles of six genes. Circles indicate decision nodes with the specified condition, classified into the signaling regulators (blue circles) and TFs (red circles). Square boxes show the number of samples and the ratio of basal-like subtype samples in the branch. For example, the top circle written as “TNF¿*P*_90.85%_” has two branches among which the right branch with 93 samples satisfy the condition while the left branch includes remaining 923 samples.

We found that the expression level of TNF gene was the most important factor to distinguish basal-like to non-basal-like tumors. Group of samples having TNF expression higher than *P*_90.85%_ had 79.57% of basal-risk, which was 4.6 times larger than non-basal-like group. Then, remaining 923 samples was split into two groups by the DT model based on ELK1 expression level, which implied that ELK1 is the second most important factor. In this case, group of samples having ELK1 expression higher than *P*_85.83%_ had 51.79% basal-risk, which was 4.1 times larger than non-basal-like group. Lastly, 112 samples having ELK1 expression higher than *P*_85.83%_ were split into two groups by the DT model. The group of samples having ELK1 expression higher than *P*_93.11%_ had 64.15% of basal-risk, which was 1.6 times larger than non-basal-like group. In summary, the DT model indicates TNF is the most important and dominant factor in deciding whether a sample is basal-like or not. ELK1 was the second important factor and the most dominant factor to samples having TNF expression lower than *P*_90.85%_.

### 3.3.2 Clinical implication of crosstalk between TNF and ELK1 and the contribution of the proposed algorithm

Results from the DT model analysis implies that the patient group with high TNF expression is riskier to be basal-like subtype. When TNF expression is not that high, patient groups with high ELK1 expression still has higher risk to be basal-like (Figure 4). To investigate that the risk of basal-like cancer predictable from the expression of TNF and ELK1 is associated with actual clinical outcome, we performed a survival analysis by drawing Kaplan-Meier plot with measuring log-rank test p-value, using TCGA-BRCA data (Supplementary Figure S11). First, we tested survival difference between TNF-high (¿*P*_90.85%_) and TNF-low group (¡=*P*_90.85%_), whose log-rank test p-value was 0.034. The result showed that the group of patients having TNF expression higher than *P*_90.85%_ had poorer survival outcome, which was consistent with the DT analysis. However, we were not able to find significant survival differences between the three patient groups separated by ELK1 expression at *P*_85.83%_ and *P*_93.11%_ under TNF *≤ P*_90.85%_, whose log-rank test p-value was 0.91. Therefore, we conjecture that TNF expression has a significant association with patient survival as well as basal-risk while the prognostic effect of ELK1 is not conclusive despite its association with basal-risk.

According to Spearman’s correlation between expressions of TNF and ELK1, two genes have significant positive correlation (p-value: 4.83E-7). The smoothed plot in Supplementary Figure S11 also shows that TNF expression is strongly associated with ELK1 expression. Moreover, the signaling path deduced by Pathway Logic also indicates that ELK1 is associated with TNF (Figure 3). However, the survival analysis result implies that ELK1 expression level do not have prognostic power. A possible explanation is that ELK1 indirectly increases basal-risk by activating TNF but does not have a direct effect on malignancy by itself. This hypothesis is in part supported by current research findings in that TNF has been considered as a molecular target in breast cancer [3] but no relevant evidence is found for ELK1.

### 3.4 Performance comparison with existing algorithms

To measure the performance of our proposed system, we analysed the same TCGA breast cancer dataset with two different algorithms: ARACNE and Clipper.

ARACNE [8] was selected as a representative of the reverse engineering algorithm. In the regulatory network predicted by ARACNE, we first filtered the edges with three criteria: 1) both genes in the edge were differentially expressed between cancer subtype and normal samples, 2) the likelihood score of the edge from ARACNE was larger than 0.5 and 3) the type of the edge (activator, suppressor) should be consistent with gene expression information (up- or down-regulated, respectively). Then, we mapped genes in the remaining edges to EGF, TNF, TGFB1 pathways in KEGG [12] database. The results are shown in Supplementary Figures S12-S14. Blue squares denote TFs and red squares denote mapped genes. As shown in the figure, most of the mapped genes were scattered and segmented in the pathways. Some of connected paths in KEGG diagram (e.g. top path in TGFB1, Luminal B subtype) were actually not connected in the ARACNE network. Clipper [14] is an algorithm for detecting signaling paths specific to a specific phenotype. Gene expressions of each cancer subtype and normal samples were analysed by Clipper to identify subtype-specific signal paths. As shown in Supplementary Figures S15-S16, signaling paths detected by Clipper were similar to each other for all subtypes, thus they were not informative for the subtype analysis of cancer.

These experiments showed that signaling pathways are unlikely to be detected from the gene expression difference only, which supports the motivation of our study.

## 4 Discussion

We proposed a new computational framework to integrate pathway models of PL, gene expression data and transcriptional network to explore subtype-specific biological mechanisms in breast cancer. Models in PL are constructed with detailed descriptions including cellular locations, interactions and PTMs of the molecules. In addition, introducing a transcriptional network into the PL model can lead to the subsequent global changes in a cell such as pathway crosstalk.

Analysing the TCGA breast cancer dataset, we derived paths of signal transduction from key receptors (TNFR, TGFBR1 and EGFR) to key TF regulators (RELA, ATF2, SMAD3 and ELK1) and to downstream genes targeted by TF, specifically detected in a certain subtype. From the basal-specific signaling paths and TFs reported, we derived a pathway crosstalk between three signaling pathways mediated by three TFs, which could provide a novel insight on aggressive breast cancer subtypes. To support the hypothesis that the pathway crosstalk with six genes is the characteristic of basal-like breast cancer, we performed decision tree and survival analysis. The analysis results indicate TNF has strong association with basal-like subtype and bad prognosis, which is supported by the literature, showing the predictive power of the proposed model. ARACNE, a well-known reverse engineering system, is also applied to the same dataset for performance comparison. Regulatory trees of TFs from ARACNE mapped to TNF, TGFB1 and EGF signaling pathways are scattered and shorter than the paths from our methods, showing less efficiency to characterise subtype-specific signaling path.

