## Supplementary for "Logic-based Analysis of Gene Expression Data Predicts Pathway Crosstalk between TNF, TGFB1 and EGF in Basal-like Breast Cancer"

### Supplementary Information for: Logic-based Analysis of Gene Expression Identifies Potential Pathway Crosstalk between TNF, TGFB1 and EGF in Basal-like Breast Cancer

Kyuri Jo, Beatriz Santos Buitrago, Minsu Kim, Sungmin Rhee, Carolyn Talcott, Sun Kim

**2 Supplementary Note: Subtype-specific path in TGFB1 pathway to SMAD3** **7**

**3 Supplementary figures** **9**

**4 Supplementary table** **23**

### 1 Supplementary Note: Pathway Logic dishes — TNF, TGFB1 and EGF

#### 1.1 Tumor Necrosis Factor signaling pathway

Tumor necrosis factor (TNF)is a cell signaling protein that belongs to a superfamily of cytokines. It mediates in a wide variety of activities, such asapoptoticcell death andimmunity. It has been implicated in the pathogenesis of a wide spectrum of human diseases, including cancer. The initial step in TNF signaling involves the binding of the TNF trimer to the extracellular domain of TNFR1 and the release of the inhibitory protein silencer of death domains (SODD). The resulting aggregated TNFR1 ICD is recognized by the adaptor protein TRADD, which recruits additional adaptor proteins RIP, TRAF2, and FADD. These latter proteins recruit key enzymes to TNFR1 that are responsible for initiating signaling events. Three pathways can be initiated: (Wajant *et al.*, 2003; Chen and Goeddel, 2002) NF-kappa-B, MAPKpathways and induction of death signaling.

**TNF dish and rewrite rules**

An initial state or *dish* (called `TnfDish`) with several locations and elements is defined:

- the outside (location tag `XOut`) which contains the tumor necrosis factor (`Tnf`);
- the `TnfR1C` location which contains the tumor necrosis factor receptor 1 (`TnfR1`);
- the `EgfRC` location which contains the epidermal growth factor receptor (`EgfR`);
- the `CLo` location, which contains the elements stuck to the outside of the plasma membrane, is empty;
- the membrane (location tag `CLm`) contains protein `Muc1`;
- the inside of the membrane (location tag `CLi`) contains several proteins `Cltc`, `Pld1`, and `Pld2`;
- the cytoplasm (location tag `CLc`) contains protein `Gsk3s` activated and some proteins `Akt1`, `Ask1`, `Azi2`, `Ciap1`, etc.;
- the nucleus (location tag `NUc`) contains several genes (e.g., `CcL2-gene`, `CcL5-gene`, `Cezanne-gene`, etc.) and proteins (e.g., `Atf1`, `Egr1`, and so on).

In Maude syntax, this *dish* (called `TnfDish`) is expressed by the following equation:

```
eq TnfDish =
PD({XOut | Tnf} {TnfR1C | TnfR1} {EgfRC | EgfR}
{CLo | empty} {CLm | Muc1} {CLi | Cltc Pld1 Pld2}
{CLc | [Gsk3s - act] Akt1 Ask1 Atf2 Azi2 Ciap1 Csnk1a1 Eif4e Eif4ebp1 Erks
     Ikba Ikbb Ikk1 Ikk2 Irak1 Itch Jnks Map4k5 Mapkapk2 Mek1 Mekk1 Mekk2
     Mekk3 Mkk4 Mkk7 Mlk3 Mlst8 Mnk1 Mtor Nemo P38s Pin1 Pkci Pkcz Raptor
     Rela Rela Ripk1 Rnf11 Rsk2 S6k1 Stat1 Syk Tab1 Tab2 Tab3 Tak1 Tank
     Tax1bp1 Tbk1 Tnfaip3 Tpl2 Tradd Traf2 Tsc1 Ubc13}
```

```
{NUc | Atf1 CcL2-gene CcL5-gene Cezanne-gene Creb1 CxcL1-gene CxcL2-gene
     CxcL10-gene Ep300 Egr1 Egr1-gene Eselectin-gene Fos-gene Cd54-gene Ikba
     -gene IL1b-gene IL6-gene IL8-gene Irf1-gene Jun Junb-gene Msk1 TLR2-
     gene ProTnf-gene Tnfaip3-gene}) .
```

The full set of rules in the TNF pathway logic model can be found at `http://stella.csl.sri.com/~pl/Evidence/STM7/rules/TnfRules.maude`. Datums for each rule in the TNF model can be found at `http://stella.csl.sri.com/~pl/Evidence/STM7/evidence/Tnf-Evidence`.

##### 1.1.1 Rewrite rule `107.TnfR1.by.Tnf`

Our rewrite rule 107 establishes: *In the presence of tumor necrosis factor* `Tnf` *in the outside of the cell (*`XOut`*), the receptor* `TnfR1` *binds to* `Tnf` *(*`TnfR1 : Tnf`*).*

In Maude syntax, this signaling process is expressed by the following rewrite rule:

```
rl[107.TnfR1.by.Tnf]:
{XOut | xout Tnf} {TnfR1C | tnfr1c TnfR1}
=> {XOut | xout} {TnfR1C | tnfr1c (TnfR1 : Tnf) } .
```

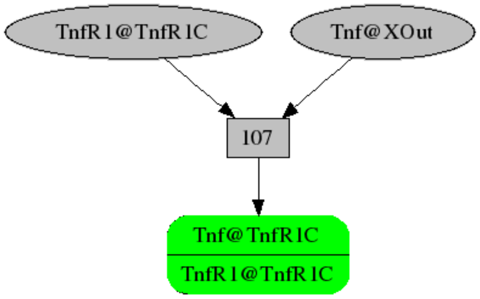

Figure N1: Visual representation of rule `[107.TnfR1.by.Tnf]` using Pathway Logic Assistant.

#### 1.2 Transforming Growth Factor Beta 1 signaling pathway

Transforming growth factor beta 1 (TGF-beta1) is the prototypic member of a large family of structurally related pleiotropic-secreted cytokines. The TGF-beta1 signaling pathway is involved in many cellular processes such as growth, proliferation, differentiation and apoptosis. The initial step in TGF-beta1 signaling involves the binding of extracellular TGF-beta1 protein to the TGF-beta type II receptor on the cell surface. This causes the recruitment and dimerization of type II receptors. These receptors recruit and phosphorylate the TGF-beta type I receptors, which phosphorylate the receptor-regulated SMAD (SMAD2 and SMAD3) presented by the SMAD anchor for receptor activation. The phosphorylated receptor-regulated

SMAD form heterologous complexes with the common-mediator SMAD (SMAD4) and subsequently translocate into the nucleus, where they interact with other transcription factors to regulate the expression of target genes (Su *et al.*, 2015; Massagué, 1998)

**TGFB1 dish and rewrite rules**

An initial state or *dish* (called `Tgfb1Dish`) with several locations and elements is defined:

- the outside (location tag `XOut`) which contains the transforming growth factor beta 1 (`Tgfb1`);
- the `Tgfb1RC` location which contains the transforming growth factor beta receptor I (`TgfbR1`) and II (`TgfbR2`);
- the `CLo` location, which contains the elements stuck to the outside of the plasma membrane, is empty;
- the membrane (location tag `CLm`) is empty as well;
- the inside of the membrane (location tag `CLi`) contains three proteins bound to GDP `Cdc42`, `Hras`, and `Rac1`;
- the cytoplasm (location tag `CLc`) contains proteins `Abl1`, `Akt1`, `Atf2`, `Erks`, etc.;
- the nucleus (location tag `NUc`) contains several genes (e.g., `Smad7`, `Tgfb1`, `Cst6`, etc.) and proteins (e.g., `Ctdsp1`, `Ets1`, and so on).

In Maude syntax, this *dish* (called `Tgfb1Dish`) is expressed by the following equation:

```
eq Tgfb1Dish =
PD({XOut | Tgfb1} {Tgfb1RC | TgfbR1 TgfbR2} {CLo | empty}
{CLm | empty} {CLi | [Cdc42 - GDP] [Hras - GDP] [Rac1 - GDP]}
{CLc | Abl1 Akt1 Atf2 Erks Fak1 Jnks Mekk1 Mlk3 P38s Pak2 Pml Smad2 Smad3
     Smad4 Smurf1 Smurf2 Tab1 Tab2 Tab3 Tak1 Traf6 Zfyve16}
{NUc | Ctdsp1 Ets1 Smad7 Cdc6-gene Cdkn1a-gene Cdkn2b-gene Col1a1-gene
     Col3a1-gene Ctgf-gene Fn1-gene Mmp2-gene Pai1-gene Smad6-gene Smad7-
     gene Tgfb1-gene Timp1-gene Cst6-gene Dst-gene Mmp9-gene Mylk-gene Pthlh
     -gene Gfi1-gene Csrp2-gene RoRc-gene}) .
```

The full set of rules in the TGFB1 pathway logic model can be found at `http://stella.csl.sri.com/~pl/Evidence/STM7/rules/Tgfb1Rules.maude`. Datums for each rule in the TGFB1 model can be found at `http://stella.csl.sri.com/~pl/Evidence/STM7/evidence/Tgfb1-evidence`.

##### 1.2.1 Rewrite rule `931.TgfbR1.TgfbR2.by.Tgfb1`

Our rewrite rule 931 establishes: *In the presence of transforming growth factor beta receptor I* ***Tgfb1*** *in the outside of the cell (****XOut****), the receptors* ***TgfbR1*** *and* ***TgfbR2*** *get activated (* ***TgfbR1-act*** *and* ***TgfbR2-act****) and bound between them and to Tgfb1 (* ***([TgfbR1 - act] : [TgfbR2 - act] : Tgfb1)*** *).*

In Maude syntax, this signaling process is expressed by the following rewrite rule:

```
rl[931.TgfbR1.TgfbR2.by.Tgfb1]:
{XOut | xout Tgfb1 } {Tgfb1RC | tgfb1rc TgfbR1 TgfbR2 }
=>
{XOut | xout }
{Tgfb1RC | tgfb1rc ([TgfbR1 - act] : [TgfbR2 - act] : Tgfb1) } .
```

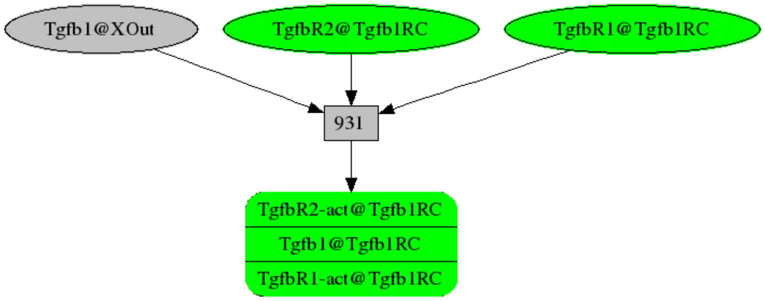

Figure N2: Visual representation of rule `[931.TgfbR1.TgfbR2.by.Tgfb1]` using Pathway Logic Assistant.

#### 1.3 Epidermal Growth Factor signaling pathway

The ErbB family of the receptor tyrosine kinases contains the epidermal growth factor receptor (EGFR) (Herbst, 2004; Normanno *et al.*, 2006). They couple the binding of the extracellular growth factor ligands to intracellular signaling pathways that control various biologic responses such as proliferation, differentiation, cell motility and survival (Engelman, 2009; Krasinskas, 2011; Marmor *et al.*, 2004; Nicholson and Anderson, 2002; Sasaki *et al.*, 2013; Suhardja and Hoffman, 2003). Three major steps are involved in the activation of EGFR-dependent intracellular signaling (Katzel *et al.*, 2009): (a) the binding of a receptor-specific ligand takes place in the extracellular portion of the EGFR or of one of the EGFR-related receptors, (b) the formation of a functionally active EGFR-EGFR dimer or an heterodimer causes the ATP-dependent phosphorylation of specific tyrosine residues in the EGFR intracellular domain, and (c) this phosphorylation triggers a complex program of intracellular signals to the cytoplasm and then to the nucleus.

EGFR activates two major intracellular pathways: (i) the RAS-RAF-MEK-MAPK pathway, which controls gene transcription, cell-cycle progression from the

G1 phase to the S phase, and cell proliferation; and (ii) the PI3K-Akt pathway, which activates a cascade of anti-apoptotic and prosurvival signals: bFGF (basic fibroblast growth factor), HB-EGF (heparin-binding EGF), MAPK (mitogen-activated protein kinase), PI3K (phosphatidylinositol 3,4,5-kinase), TGF-$\alpha$ (transforming growth factor alpha), and VEGF (vascular endothelial growth factor).

The binding between EGFR and ligand triggers downstream intracellular signaling pathways. Some of them include the PI3K-Akt prosurvival, STAT transcription, and RAS-RAF-MEK proliferation pathways (Marmiroli *et al.*, 2015). The RAS-RAF-MEK and PI3K-Akt pathways are mostly activated by the anaplastic lymphoma kinase (ALK) fusion proteins. Cell proliferation, cell motility and carcinogenesis is impulsed by the amplification of the EGFR and ALK signaling pathways.

**EGF dish and rewrite rules**

An initial state or *dish* (called `EgfDish`) with several locations and elements is defined:

- the outside (location tag `XOut`) which contains the epidermal growth factor (`Egf`);
- the `EgfRC` location which contains the epidermal growth factor receptor (`EgfR`);
- the `CLo` location, which contains the elements stuck to the outside of the plasma membrane, is empty;
- the membrane (location tag `CLm`) contains proteins `Erbb2`, `Pag1`, and `Plscr1`;
- the inside of the membrane (location tag `CLi`) contains several proteins binding to guanosine diphosphate `GDP`: `Cdc42`, `Hras`, `Kras`, etc.; and some other proteins: `Gnai1`, `Gnai3`, `Pld1`, etc. (see the full code below);
- the cytoplasm (location tag `CLc`) contains enzyme `Pi3k` and some proteins `Abl1`, `Akt1`, `Araf`, `ArhGap5`, etc.;
- the nucleus (location tag `NUc`) contains several proteins (e.g., `Atf1`, `Creb1`, `Elk1`, and so on).

In Maude syntax, this *dish* (called `EgfDish`) is expressed by the following equation:

```
eq EgfDish =
PD({XOut | Egf} {EgfRC | EgfR}{CLo | empty} {CLm | Erbb2 Pag1 Plscr1}
{CLi | [Cdc42 - GDP] [Hras - GDP] [Kras - GDP] [Nras - GDP] Gnai1 Gnai3 [
     Rac1 - GDP] [Rala - GDP] [Ralb - GDP] [Rap1a - GDP] Pld1 Pld2 [Rap2b -
     GDP] [Rit1 - GDP] Src}
{CLc | (Mlst8 : Mtor : Raptor) [Gsk3s - act] Abl1 Akt1 Araf ArhGap5 ArhGef4
     ArhGef7 Atf2 Bmx Braf Cbl Cblb Cin85 Crk CrkL Csk Dbl Dok1 Dok2
     Eif4ebp1 EndA1 Eps8 Eps15 Erk5 Erks Fak1 Fak2 Flna Freud1 Gab1 Gab2
     Git1 Grb2 Hpk1 Ipo7 IqGap1 Jak1 Jak2 Jnks Lkb1 Mapkapk2 Mek1 Mek5 Mekk1
      Mekk2 Mekk3 Mlk3 Nckipsd P38s Pdpk1 Pi3k Pkcd Pkcz Plcg1 Plce1 Ptk6
     Pxn Raf1 RalGds RapGef1 Rasa1 RasGrp3 Rela Rictor Rin1 Rps6 Rsk1 Rsk2
     Rsk3 S6k1 Sh2d3a Sh2d3c Shc1 Shoc2 Shp2 Sin1 Smad3 Sos1 Stat1 Stat3
     Stat5a Stat5b Tnk2 Tns3 Ube2l3 Vav1 Vav2 Vav3 Ywhaz }
```

```
{NUc | Atf1 Creb1 Egr1-gene Elk1 Fos HistH3 Irf1-gene Jun Jun-gene Junb-gene
     Msk1 Msk2 Myc Myc-gene Nos2a-gene Socs3-gene Tbp-gene} ) .
```

Rewrite rules detail the behavior of cell components depending on biological contexts and modification states. Each rule represents an action in a biological process such as intra/inter cellular signaling reactions or metabolic reactions. Pathway Logic contains a set of transition rules, derived from curated experimental findings. They provide an explanation of how a signal propagates in response to an EGF stimulus. The EGF pathway logic model contains a total of 129 rules and 2281 datums. The full set of rules in the EGF pathway logic model can be found at `http://stella.csl.sri.com/~pl/Evidence/STM7/rules/EgfRules.maude`. The experimental evidence for each rule is supplied in datum form. Each datum represents a result from a experiment published in a refereed journal. Datums for each rule in the EGF model can be found at `http://stella.csl.sri.com/~pl/Evidence/STM7/evidence/Egf-Evidence`.

##### 1.3.1 Rewrite rule `001.EgfR.irt.Egf`

Here we describe rule 001, directly sourced from the literature. Yarden *et al.* (Yarden and Sliwkowski, 2001) determine that when EGF and its relatives bind the ErbB family of receptors, they trigger a network of signaling pathways, conclude with responses ranging from cell division to death, motility to adhesion.

Rewrite rule 001 establishes: *In the presence of epidermal growth factor* ***`Egf`*** *in the outside of the cell (`XOut`), the receptor* ***`EgfR`*** *is phosphorylated on tyrosine (`[EgfR - Yphos]`) and binds to protein EGF (`Egf`)* (Elenius *et al.*, 1997; Yarden and Sliwkowski, 2001). In Maude syntax, this signaling process is expressed by the following rewrite rule:

```
rl[001.EgfR.irt.Egf]:
{XOut | xout Egf}  {EgfRC | egfrc EgfR}
=>  {XOut | xout}  {EgfRC | egfrc ([EgfR - Yphos] : Egf)} .
```

Fig. N3 gives a Petri net representation of the aforementioned rule using the Pathway Logic Assistant. An oval represents a component (e.g., gene, protein, etc.) participating in a reaction. A rectangle illustrates a reaction rule with a label which represents its shortened identifier in the knowledge base. A solid arrow from an occurrence oval to a rule indicates that the occurrence is a reactant. A solid arrow from a rule to an occurrence oval indicates that the occurrence is a product.

### 2 Supplementary Note: Subtype-specific path in TGFB1 pathway to SMAD3

SMAD3-path was captured in TGFB1 pathway because its transcriptional target genes are luminal B-specific DEGs. SMAD3-path consists of 3 components, including Tab1/Tak1, JNK, and SMAD3. Here, the first two components (Tab1/Tak1

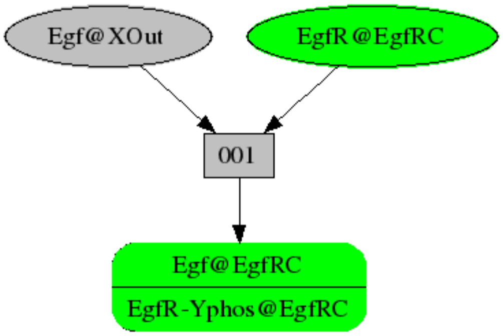

Figure N3: Visual representation of rule `[001.EgfR.irt.Egf]` using Pathway Logic Assistant.

and JNK) are cascading kinases transmitting TGFB1 signal into nucleus to activate SMAD3, which subsequently induce transcription of its target genes (Figure N4).

According to our findings summarized in Table N1, SMAD3 is over-expressed specifically in luminal B patients (and less significantly in luminal A), while HER2-enriched and basal-like patients show rather down-regulated patterns. Since SMAD3 showed up-regulated patterns only in the good prognostic group (i.e. luminal A and B) and showed down-regulated patterns in the bad prognostic group (i.e. HER2-enriched, basal-like), we speculate that the up-regulation of SMAD3 has a significant effect on the prognosis of each patient. What is interesting here is that SMAD3 was a downstream target that was supposed to be activated by TGFB1 signaling. So, if SMAD3 is active, it might be the result of activation of the TGFB1 signaling pathway. Therefore, unlike the case of SMAD7-path, activation of the TGFB1 pathway appears to play a tumor-suppressive role in this context. These seemingly conflicting results indicate that there might be other factors that determine whether TGFB1 pathway acts as a suppressor or inducer of cancer. We suspect that epigenomic factors are likely to be responsible for this dual nature, since there are studies suggesting that DNA methylation governs the SMAD3's role in cancer patients (Vidakovic *et al.*, 2015; Javle *et al.*, 2014).

Table N1: Gene expression pattern in SMAD3-path. Subtypes are abbreviated with B (basal-like), H (HER2-enriched), LA (luminal A), LB (luminal B), O (other cancer subtypes) and N (normal). Differential expression between subtypes is described based on the left-side subtype.

| P<0.05 | LB vs. N | LB vs. O | B vs. N | H vs. N | LA vs. N |
|---|---|---|---|---|---|
| MAP3K7 (Tak1) | - | - | UP | - | - |
| TAB1 | - | - | - | - | - |
| MAPK8 (JNK) | - | - | - | DOWN | - |
| MAPK9 (JNK) | UP | UP | DOWN | - | - |
| MAPK10 (JNK) | DOWN | - | DOWN | - | - |
| SMAD3 | UP | UP | DOWN | DOWN | UP |

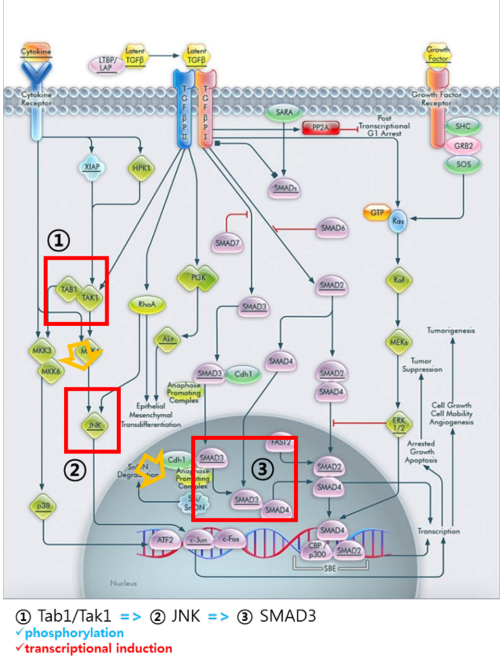

Figure N4: TGFB1 pathway marked with SMAD3-path (Source: Thermo Fisher Scientific (2011)).

### 3 Supplementary figures

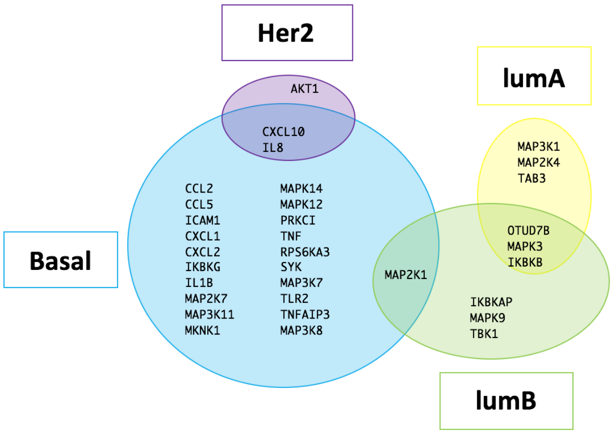

Figure S1: Venn diagram showing the DEGs in the TNF Pathway Logic model.

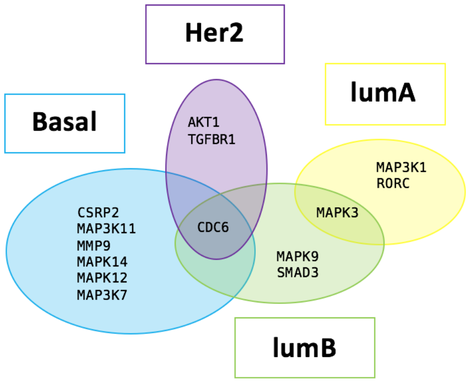

Figure S2: Venn diagram showing the DEGs in the TGFB1 Pathway Logic model.

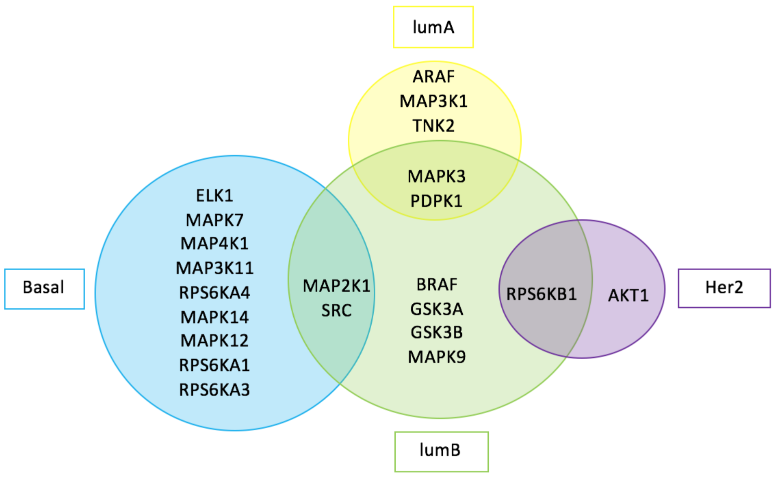

Figure S3: Venn diagram showing the DEGs in the EGF Pathway Logic model.

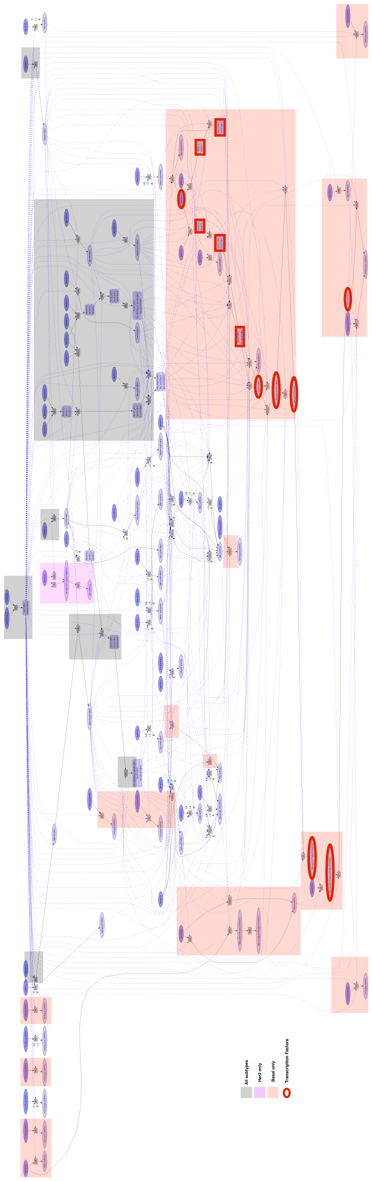

Figure S4: Transcription Factors (TFs) found in the occurrences of TNF reduced dish in Pathway Logic are marked with red ovals. Rules were labeled with the subtype that they belong to, some of which were common rule for all cancer subtypes (gray box) and some are present in only one subtype (colored box).

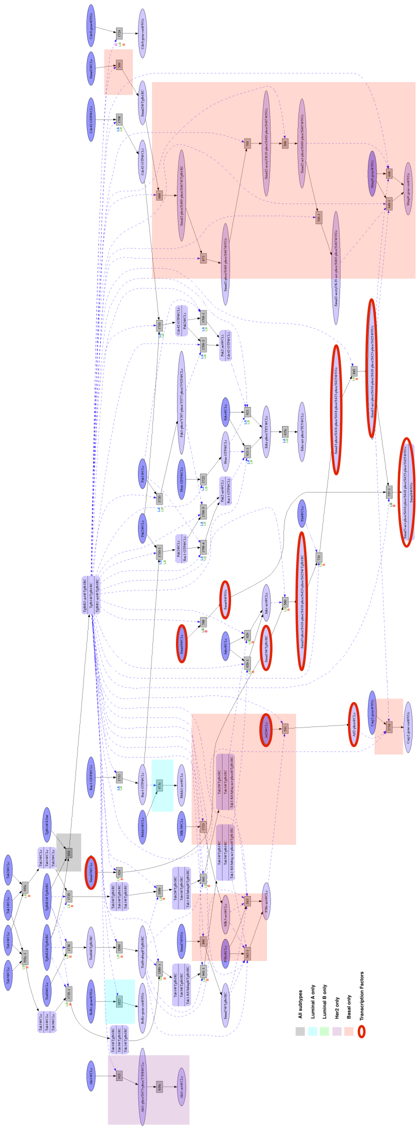

Figure S5: Transcription Factors (TFs) found in the occurrences of TGFB1 reduced dish in Pathway Logic are marked with red ovals. Rules were labeled with the subtype that they belong to, some of which were common rule for all cancer subtypes (gray box) and some are present in only one subtype (colored box).

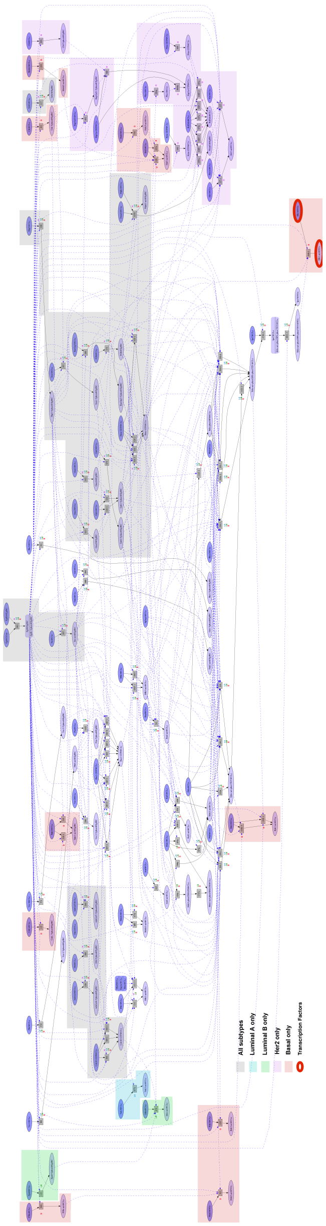

Figure S6: Transcription Factors (TFs) found in the occurrences of EGF reduced dish in Pathway Logic are marked with red ovals. Rules were labeled with the subtype that they belong to, some of which were common rule for all cancer subtypes (gray box) and some are present in only one subtype (colored box).

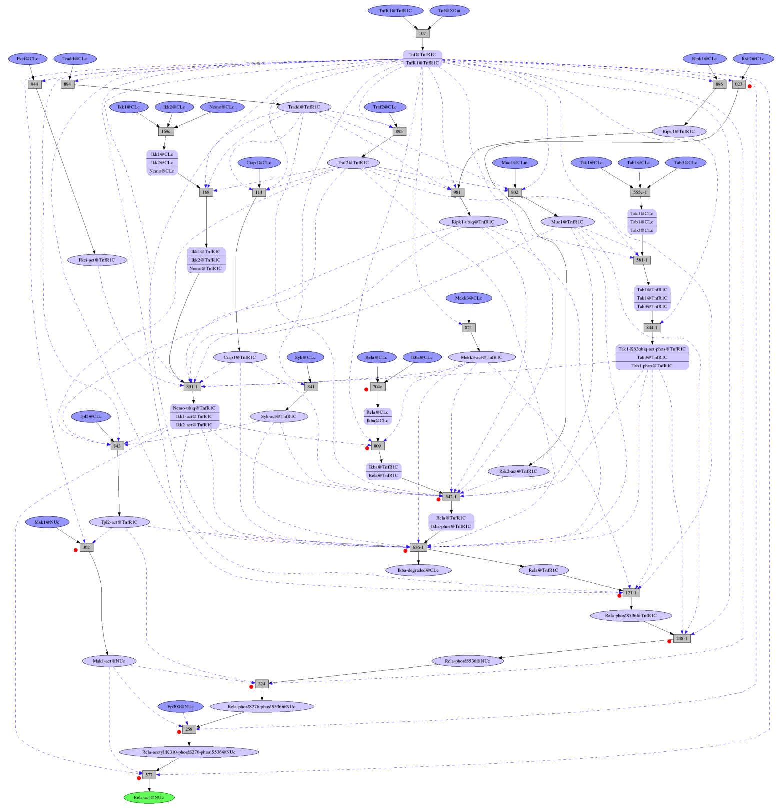

Figure S7: Subtype-specific signaling path to RELA on TNF pathway, basal-like subtype.

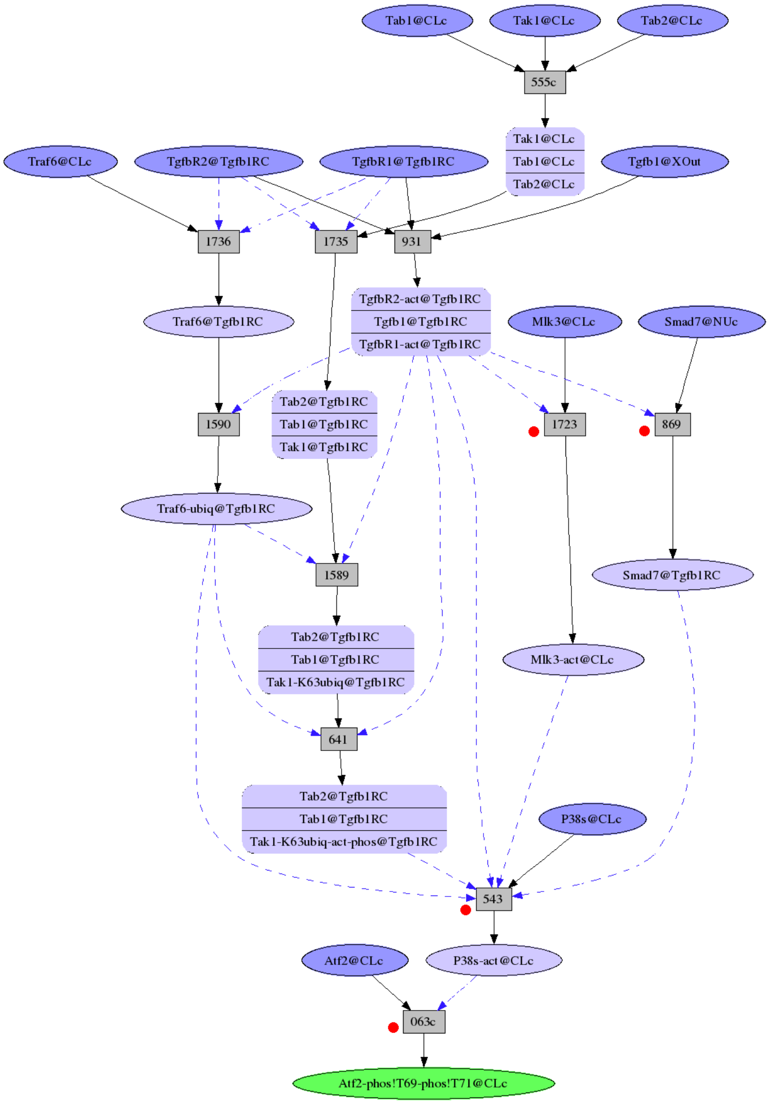

Figure S8: Subtype-specific signaling path to ATF2 on TGFB1 pathway, basal-like subtype.

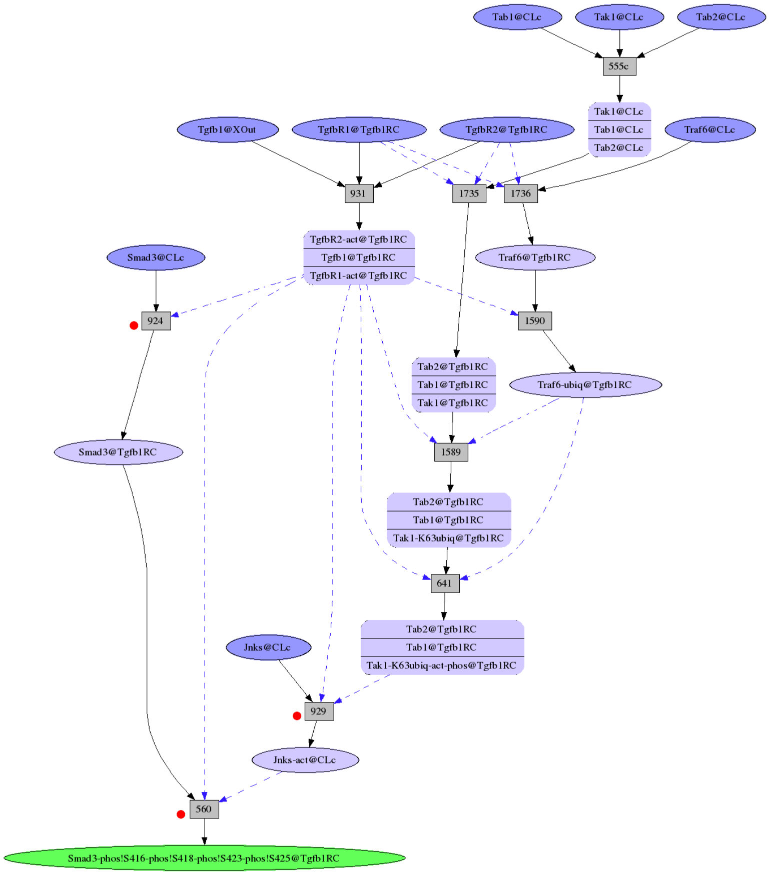

Figure S9: Subtype-specific signaling path to SMAD3 on TGFB1 pathway, luminal B subtype.

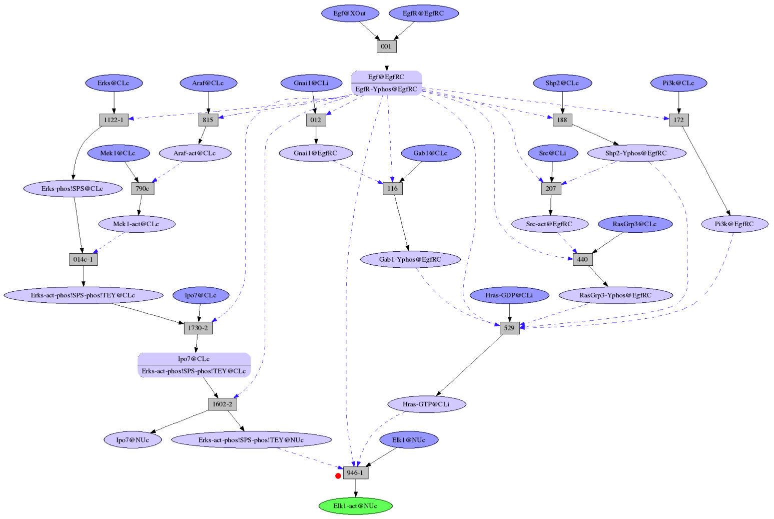

Figure S10: Subtype-specific signaling path to ELK1 on EGF pathway, basal-like subtype.

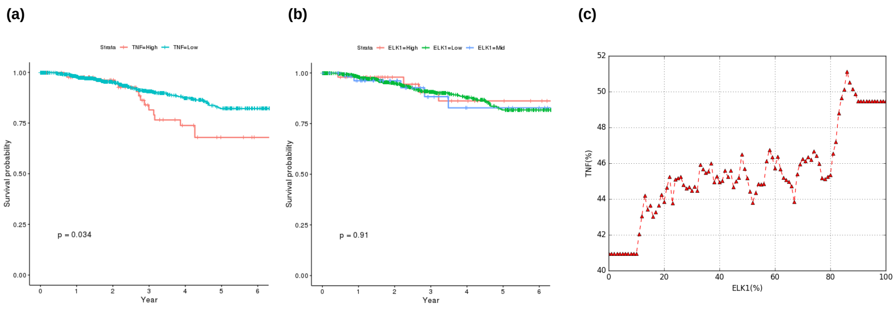

Figure S11: (a) Kaplan-Meier plot of survival analysis between TNF-high and TNF-low group. Survival analysis between two patient groups separated by DT model, where the branch condition is TNF¿$P_{90.85\%}$. TNF-high group is colored red in plot and includes 93 samples and TNF-low group is colored green and includes 923 samples. According to log-rank test (p-value 0.034), these two groups are shown to have significantly different survival outcome. (b) Kaplan-Meier plot of survival analysis between ELK-high, ELK-mid, and ELK-low group. The branch conditions are ELK1 expression $P_{93.11\%}$ and $P_{85.83\%}$. Accordingly, among 923 samples included in "TNF¡=$P_{90.85\%}$" group, 53 samples are classified as ELK1-high, 59 samples are classified as ELK1-Mid, and the remaining 811 samples are classified as ELK1-low. These three groups do not show significant survival difference (p-value 0.91). (c) Smoothed plot from ELK1 to TNF. To show the association between ELK1 and TNF, we visualize the relationship between them with a smoothed plot (see Methods). X-axis represents ELK1 expression and Y-axis represents TNF expression. The window size is 20% around x value. For example, the y-value corresponding to x-value 20% is the average TNF expression of samples included in ELK1 range (10 29%). As shown, average TNF expression is consistently elevated as ELK1 expression increases.

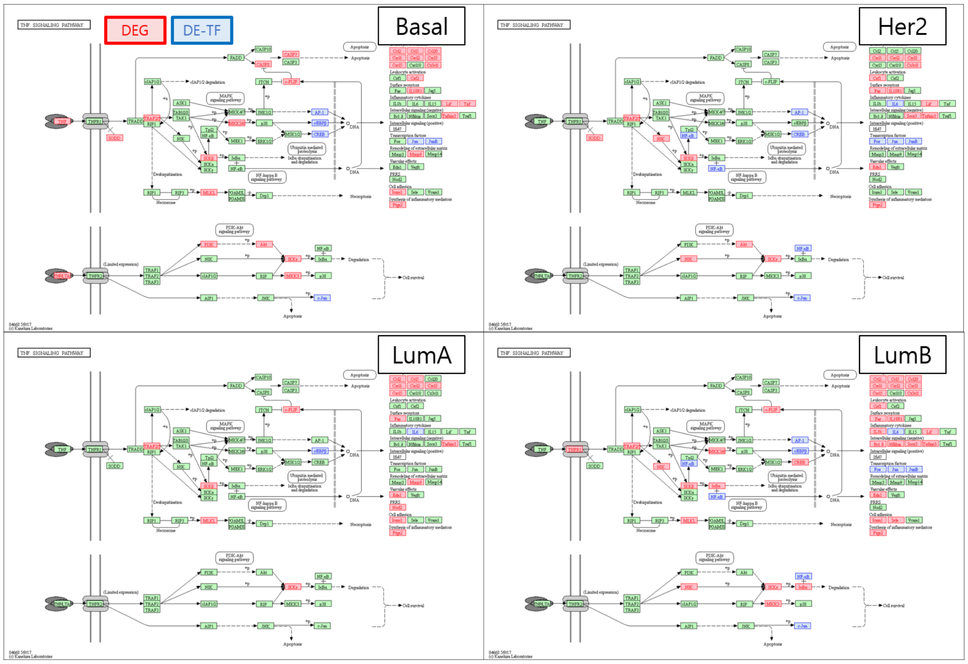

Figure S12: ARACNE results on TNF pathway.

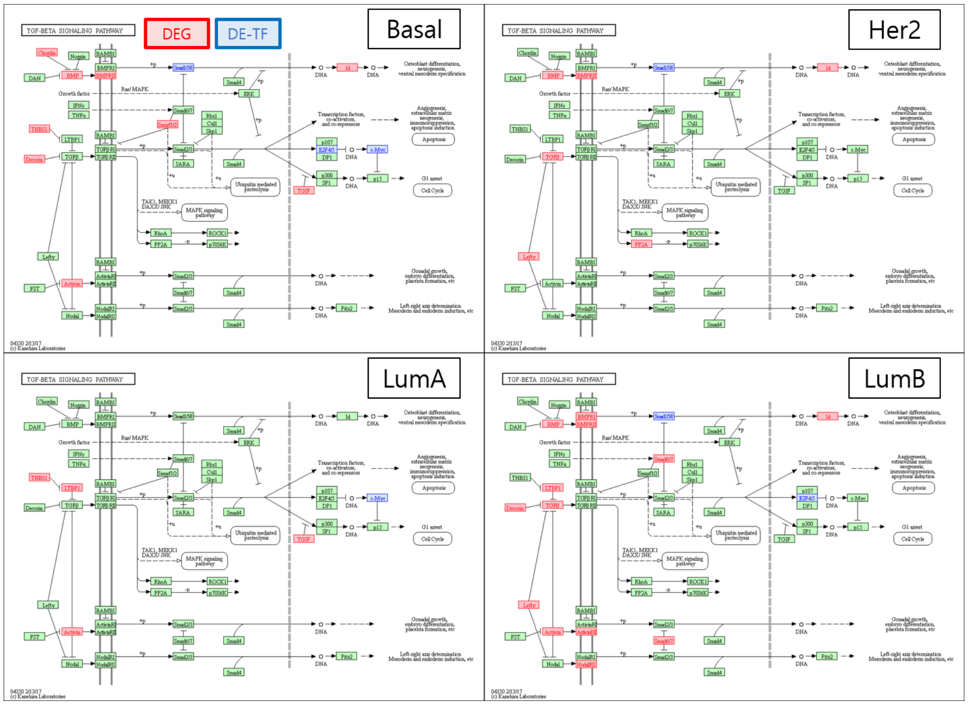

Figure S13: ARACNE results on TGFB1 pathway.

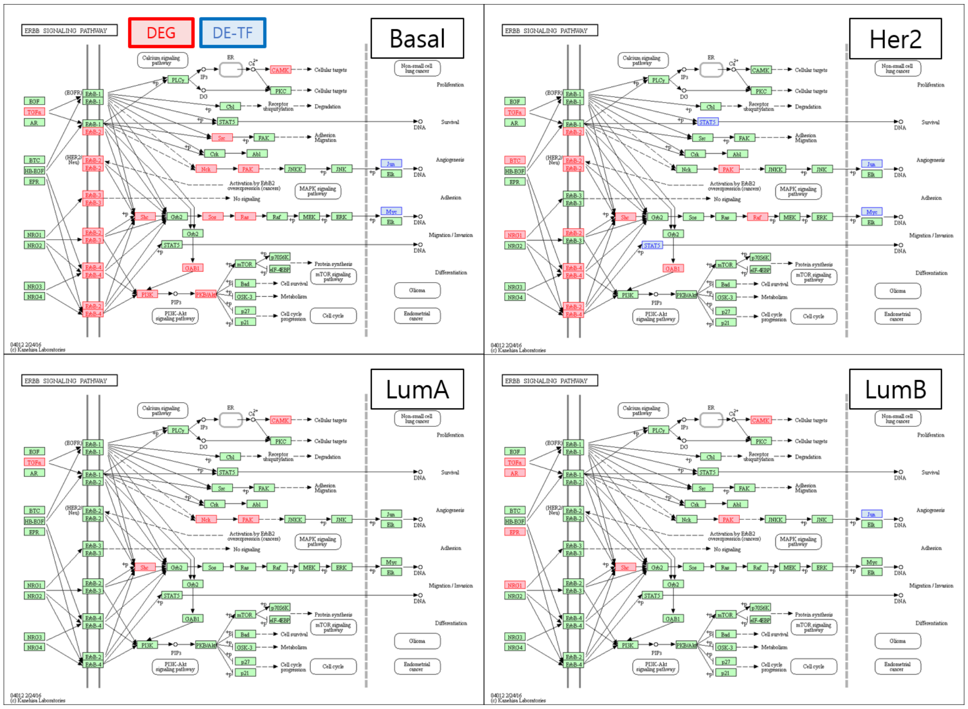

Figure S14: ARACNE results on EGF pathway.

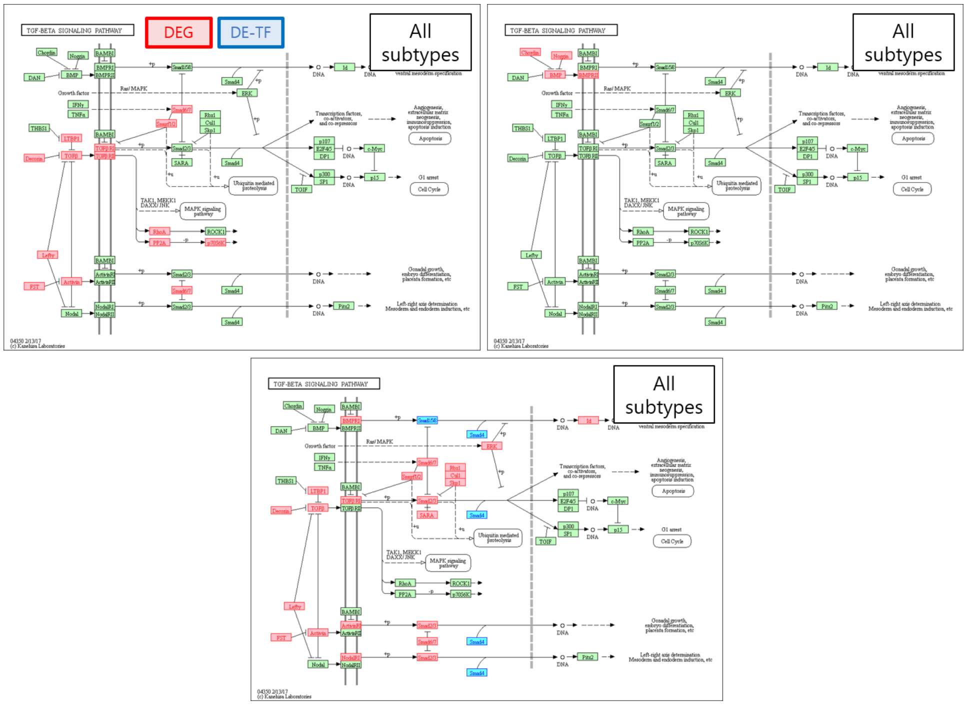

Figure S15: Clipper results on TGFB1 pathway.

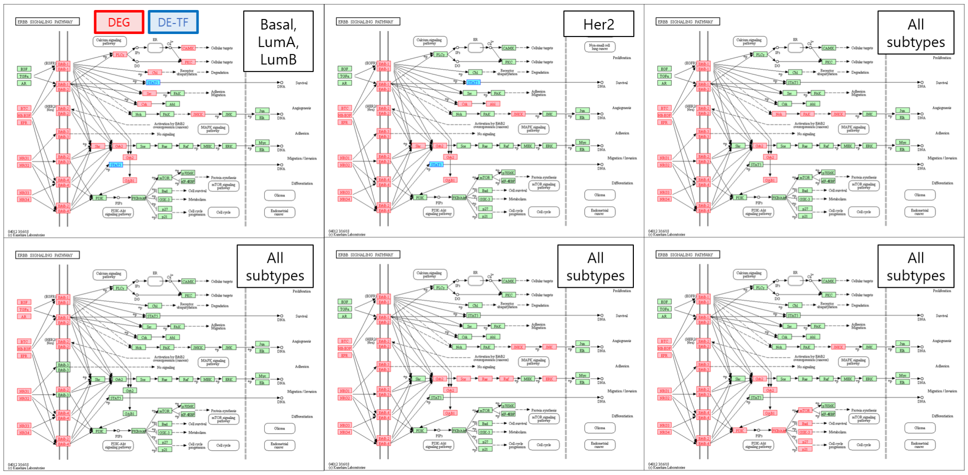

Figure S16: Clipper results on EGF pathway.

### 4 Supplementary table

Table S1: Signaling pathways connected from three upstream signaling factors (TNF, TGFB1, and EGF) to downstream crosstalk partners mediated by TFs. For each upstream pathway, TF mediators and the subtype where the crosstalk (Downstream pathway) is activated are described. Overlap indicates the number of common genes that are involved in the pathway (Pathway) and are also the differentially expressed target genes of TF (DE-TG). P-value is calculated by the enrichment analysis performed with Enrichr (Kuleshov *et al.*, 2016).

| Pathway | TF | Subtype | Downstream pathway | Overlap | Pathway | DE-TG | p-value |
|---|---|---|---|---|---|---|---|
| TNF | RELA | Basal-like | TNF signaling pathway | 21 | 110 | 174 | 2.39E-15 |
| | | | AGE-RAGE signaling pathway in diabetic complications | 17 | 101 | 174 | 9.58E-12 |
| | | | NF-kappa B signaling pathway | 16 | 93 | 174 | 2.78E-11 |
| | | | T cell receptor signaling pathway | 11 | 104 | 174 | 5.51E-06 |
| | | | MAPK signaling pathway | 17 | 255 | 174 | 1.06E-05 |
| | | | Toll-like receptor signaling pathway | 11 | 106 | 174 | 6.63E-06 |
| TGFB1 | ATF2 | Basal-like | TNF signaling pathway | 13 | 110 | 32 | 9.79E-21 |
| | | | Estrogen signaling pathway | 11 | 99 | 32 | 2.66E-17 |
| | | | Adrenergic signaling in cardiomyocytes | 11 | 148 | 32 | 2.53E-15 |
| | | | Glucagon signaling pathway | 10 | 101 | 32 | 2.87E-15 |
| | | | PI3K-Akt signaling pathway | 13 | 341 | 32 | 3.02E-14 |
| | | | AMPK signaling pathway | 10 | 124 | 32 | 2.37E-14 |
| | | | cGMP-PKG signaling pathway | 10 | 167 | 32 | 4.85E-13 |
| | | | AGE-RAGE signaling pathway in diabetic complications | 7 | 101 | 32 | 7.53E-10 |
| | | | cAMP signaling pathway | 7 | 199 | 32 | 8.44E-08 |
| | | | MAPK signaling pathway | 7 | 255 | 32 | 4.54E-07 |
| | | | HIF-1 signaling pathway | 5 | 103 | 32 | 1.44E-06 |
| | | | Jak-STAT signaling pathway | 5 | 158 | 32 | 1.17E-05 |
| | SMAD3 | Luminal B | T cell receptor signaling pathway | 13 | 104 | 180 | 2.86E-07 |
| | | | TGF-beta signaling pathway | 10 | 84 | 180 | 1.06E-05 |
| EGF | ELK1 | Basal-like | NONE | | | | |
